# An Immunoinformatics Study to Predict Epitopes in the Envelope Protein of SARS-COV-2

**DOI:** 10.1101/2020.05.26.115790

**Authors:** Renu Jakhar, S.K Gakhar

## Abstract

COVID-19 is a new viral emergent human disease caused by a novel strain of Coronavirus. This virus has caused a huge problem in the world as millions of the people are affected with this disease in the entire world. We aimed to design a peptide vaccine for COVID-19 particularly for the envelope protein using computational methods to predict epitopes inducing the immune system and can be used later to create a new peptide vaccine that could replace conventional vaccines. A total of available 370 sequences of SARS-CoV-2 were retrieved from NCBI for bioinformatics analysis using Immune Epitope Data Base (IEDB) to predict B and T cells epitopes. Then we docked the best predicted CTL epitopes with HLA alleles. CTL cell epitopes namely interacted with MHC class I alleles and we suggested them to become universal peptides based vaccine against COVID-19. Potentially continuous B cell epitopes were predicted using tools from IEDB. The Allergenicity of predicted epitopes was analyzed by AllerTOP tool and the coverage was determined throughout the worlds. We found these CTL epitopes to be T helper epitopes also. The B cell epitope, SRVKNL and T cell epitope, FLAFVVFLL were suggested to become a universal candidate for peptide-based vaccine against COVID-19. We hope to confirm our findings by adding complementary steps of both *in vitro* and *in vivo* studies to support this new universal predicted candidate.

## Introduction

As we all know the Corona virus has stopped the movements of the entire world. This virus is so deadly that it is taking lives of the more than thousands of people every day and affecting millions of people on the globe. Although the disease was first reported in the Wuhan city of China, where the virus was isolated from a patient with respiratory symptom in Dec 2019, [1, 2] later identified it by the name of COVID-19 [3]. World Health Organization (WHO) announced this disease as pandemic disease that spread from China to more than a hundred countries in the world. By May 25, 2020, the disease had already struck more than million persons of whom thousands of peoples died from COVID-19 infection majority of them were reported from China, Italy, United State of America, Britain and Spain.

Corona viruses are the large group of viruses belonging to the family Coronaviridae and the order Nidovirales that are common among animals [4]. The coronaviridae family is divided into four genera based on their genetic properties, including Alpha, Beta, Gamma and Delta corona virus genus [5]. The 2019-nCoV is enveloped positive-sense RNA, Beta corona virus with a genome of 29.9 kb [6]. They are zoonotic, transmitted from animals to humans [7]. Covid-19 affects the respiratory system (lungs and breathing tubes). Most COVID-19 patients developed severe acute respiratory illness with symptoms of fever, cough, and shortness of breath. Maximum reported cases of COVID-19 have been linked through travel to or residence in countries in this region [8, 9].

Presently there are no clinically approved vaccines available in the world for this disease. The development of new vaccine for this new emergent strain by using therapeutic and preventive approach can be readily applied to save human lives. The use of peptides or epitopes as therapeutics is a good strategy as it has advances in design, stability, and delivery [11, 12]. Moreover, there is a growing importance on the use of peptides in vaccine design by predicting immunogenic CTL, HTL and B cell epitopes from tissue-specific proteins of organisms [13, 14]. Among the structural proteins of SARS-CoV-2, the CoV envelope (E) protein is a small integral membrane protein involved in life cycle of virus. It involves in envelope formation, and some other aspects like assembly formation, budding, and pathogenesis. Thus, it is considered to be a promising target for effective COVID-19 vaccine design [15]. More importantly, T-cell-based cellular immunity is essential for cleaning SARS-CoV-2 infection because it is memory based [16, 17]. Also, the low mutation rate of the E protein or it is a highly conserved protein that can elicits both cellular immunity, and neutralizing antibody against COVID-19 is necessary for an efficient vaccine development [18, 19].

Therefore, in this study, an immunoinformatics based approach was adopted to identify a candidate epitopes against envelope protein of SARS-CoV-2 that could be appropriately activate a significant cellular, and humoral immune response [20, 21]. The aim of this study is to analyze envelope protein strains using *in silico* approaches looking for the conservancy, which is further studied to predict all potential epitopes that can be used after *in vitro* and *in vivo* confirmation as a therapeutic peptide vaccine [22, 23, 24].

## Materials and Methods

### Protein Sequence Retrieval

The protein sequence of envelope protein from severe acute respiratory syndrome coronavirus 2 isolate Indian strain (SARS-CoV-2/166/human/2020/IND) with accession no. QIA98585.1 was retrieved from the NCBI. The antigenicity of this sequence was predicted by the VaxiJen v2.0 server [25] with default parameter. In the present study, envelope protein was found to be a potential antigenic protein with good antigenicity score. A total of 370 envelope protein sequences were retrieved from the NCBI database till 12 April 2020. These 370 sequences retrieved were collected from different parts of the world; retrieved sequences and their accession numbers are listed in the supplementary file. Further, the multiple sequence alignment of envelope protein sequences was carried out through clustal W.

### Homology Modelling

Envelope protein 3D structure was obtained by Swissmodeller which uses homology detection methods to build 3D models [26]. UCSF Chimera was used to visualize and minimize the 3D structures [27], and structure validation was carried out with SAVES [28]. Homology modelling was achieved to establish conformational B cell epitope prediction and for further verification of the surface accessibility and hydrophilicity of B lymphocyte epitopes predicted, as well as to visualize all predicted T cell epitopes in the structural level.

### B-cell Epitope Prediction

B cell epitope is the portion of an immunogen, which interacts with B-lymphocytes. As a result, the B-lymphocyte is differentiated into an antibody-secreting plasma cell and the memory cell. Thus, the IEDB resource was used for analysis. Envelope protein was subjected to Bepipred linear epitope prediction [29], Emini surface accessibility [30], Kolaskar and Tongaonkar antigenicity [31], Parker hydrophilicity [32], Chou and Fasman beta turn [33] and Karplus & Schulz Flexibility Prediction [34] prediction methods in IEDB, that predict the probability of specific regions in the protein to bind to B cell receptor, being in the surface, being immunogenic, being in a hydrophilic region and being in a beta turn region, respectively. Potentially continuous B cell epitope was predicted using tool Ellipro from IEDB resource [35]. The Allergenicity of predicted epitopes was analyzed by AllerTOP Tool [36]. ToxinPred server was used to predict toxicity assessment of epitopes [37].

### MHC Class I binding predictions

T-cell epitopes were predicted by the NetCTL server [38]. The parameter was set at 50 to have the highest specificity and sensitivity of 0.94 and 0.89, respectively and all the supertypes were taken during the submission of a protein sequence. A combined algorithm of Major Histocompatibility Complex (MHC)-1 binding, Transporter of Antigenic Peptide (TAP) transport efficiency and proteasomal cleavage efficiency were used to predict the overall scores [39]. On the basis of the combined score first, five best epitopes were selected for further testing as putative epitope vaccine candidates. MHC-1 binding T cell epitope was predicted by IEDB by using the Stabilized matrix method (SMM) for each peptide [40]. Prior to prediction, all epitope lengths were set as 9mers, conserved epitopes that bind to many HLA alleles at score equal or less than 1.0 percentile rank were selected. For further analysis, alleles having IC50 less than 200 nm were selected. Overall, the higher immunogenicity of peptides shows more expected to be CTL epitopes than those having lower immunogenicity. Therefore, the IEDB immunogenicity prediction tool was used for the prediction of the immunogenicity of the candidate epitopes [41].

### Prediction of Helper T cell epitope

Analysis of peptide binding to MHC class II molecules was assessed by the IEDB MHC II prediction tool, where SMM based NetMHCIIpan 3.0 server was used [42]. It covers all HLA class II alleles including HLA-DR, HLA-DQ, and HLA-DP [43]. IC50 below 200 nM show maximum interaction potentials of HTL epitope and MHC II allele [44]. Accordingly, five top epitopes were selected. The predicted HTL epitopes were submitted to the IFN epitope server to check whether the MHCII binding epitopes had the ability to induce IFN-γ [45].

### Population Coverage Calculation

All potential MHC I and MHC II binders from envelope protein were assessed for population coverage against the whole world population that had been reported COVID-19 cases. Calculations achieved using the selected MHC-I and MHC-II interacted alleles by the IEDB population coverage calculation tool [46].

### Docking studies

Epitopes of MHC I alleles that predicted to bind with percentile rank below 0.5 were selected as the ligands, which are modeled using PEP-FOLD online peptide modeling tool [47]. The receptor MHC I allele 3D structure was obtained from the PDB server [48]. Patchdock program was used for all dockings [49]. PyMol and CHIMERA were used for visualization and determination of binding affinity and to show the suitable epitopes binding with the lowest energy.

## Results

### Retrieval of protein sequence and Antigenicity determination

The protein sequence of envelope protein from severe acute respiratory syndrome coronavirus 2 isolate Indian strain retrieved in FASTA format was screened using the VaxiJen server to predict the immunogenicity. In the present study, the QIA98585.1) was predicted to be antigenic protein based on the overall score by the Vaxijen server and this has been indicated as an immunogenic protein. A total of 370 envelope protein sequences retrieved from the NCBI database were aligned, to see the conservation of predicted epitopes. By means of IEDB analysis resource B and T cell epitopes were predicted and population coverage was calculated.

### Homology Modelling, Refinement, and Validation of E protein

Three-dimensional structure of envelope protein of the SARS-CoV-2 was modelled using the homology structure modelling tool Swissmodeller (Fig.1). This protein showed a good model with Swissmodeller by using PDB ID: 5X29 respectively as a template has more than 91% identity and 54% similarity with the query structure. These models were energy minimized by using Chimera. The Ramachandran plot and Prosa Z-score validation (Fig. 2) indicated that >86% residues in the favoured region for the modelled envelope protein.

**Fig. 1.**
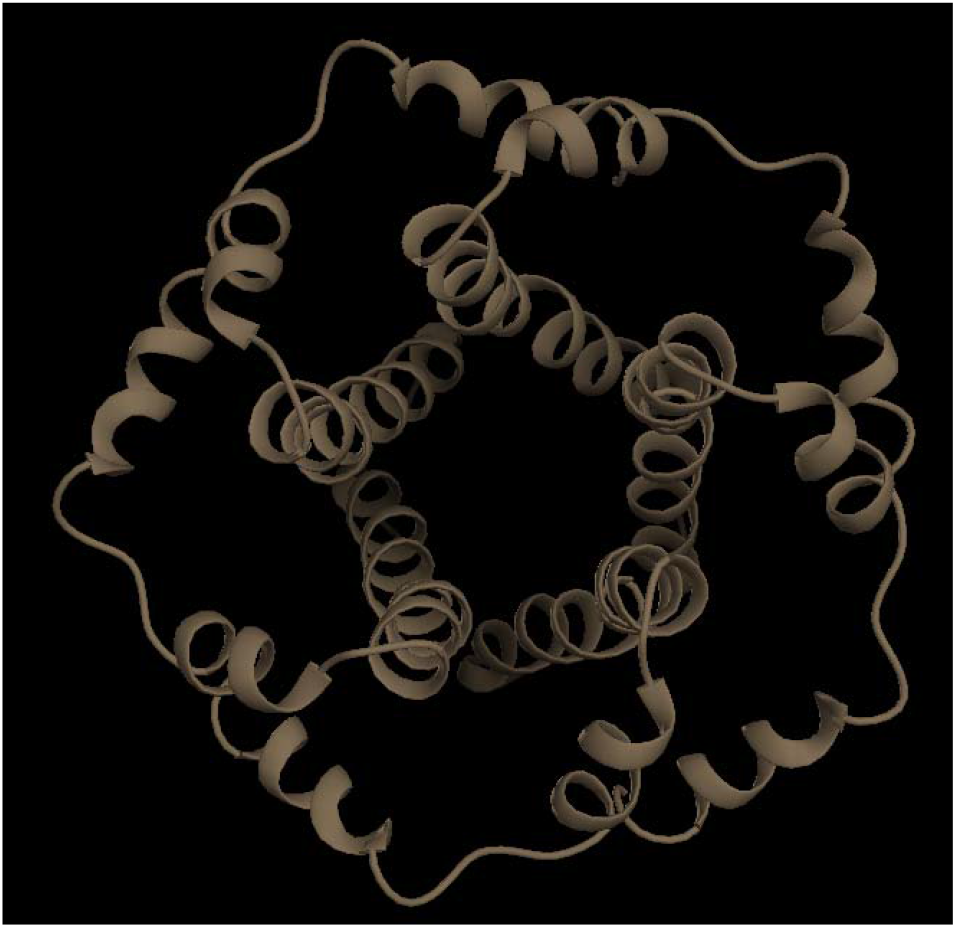
Predicted 3D structure of putative E protein of SARS-CoV-2 virus.

**Fig. 2.**
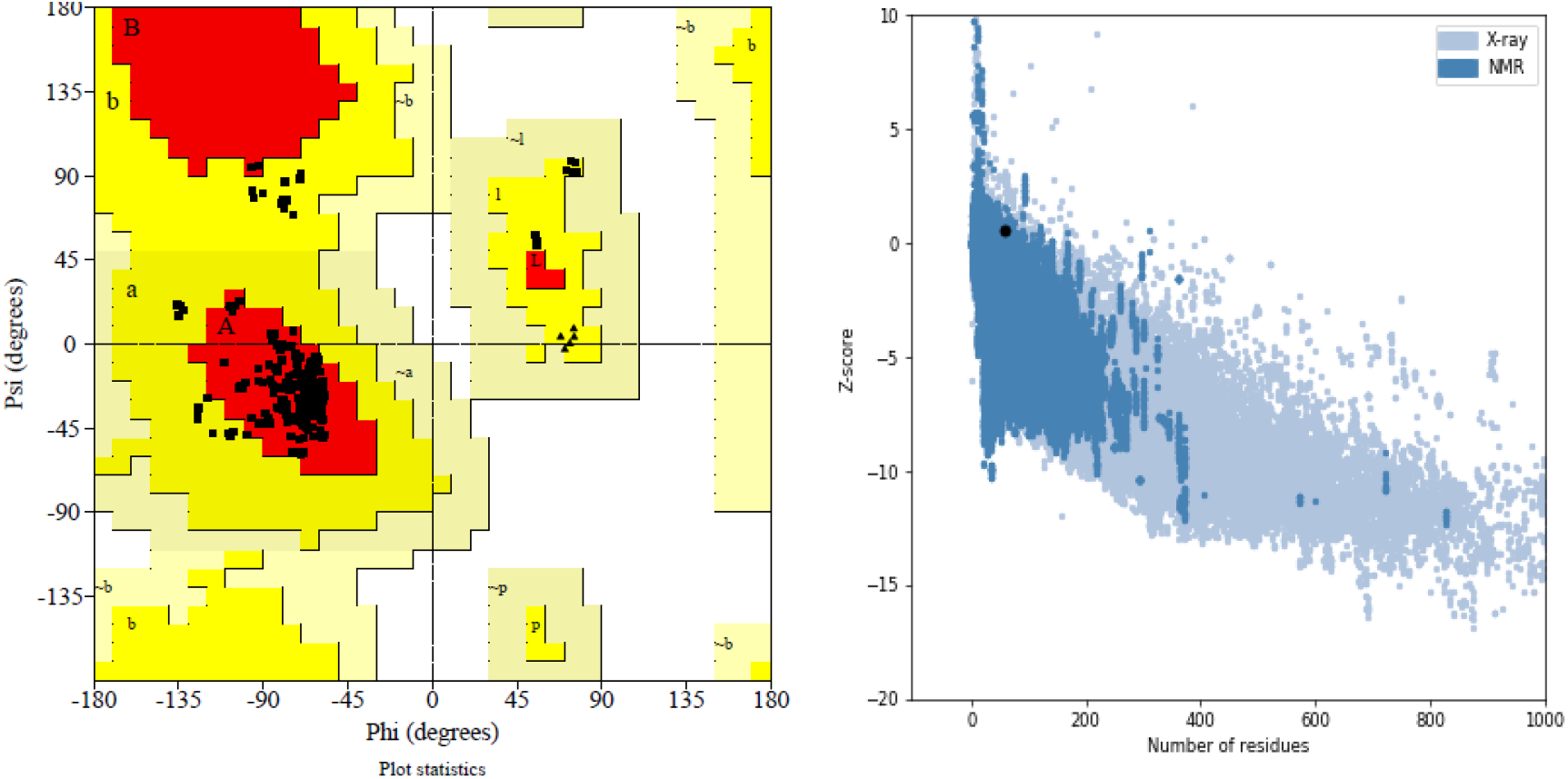
Ramachandran Plot and Prosa Z-score validation of E protein of SARS-CoV-2.

### Prediction of conformational and linear B-cell Epitope

The conformational B-cell epitopes were also obtained in five chains of envelope protein by using ElliPro. ElliPro gives the score to each output epitope, which is Protrusion Index (PI) value averaged over each epitope residue [50]. Some ellipsoids approximated the tertiary structure of the protein. The highest probability of a conformational epitope was calculated at 76% (PI score: 0.76). Residues involved in conformational epitopes, their number, location and scores are shown in Table 1, ^60^SRVKNL^65^ residues were found have highest PI score. This epitope is antigenic, nonallergic, nontoxin, and conserved in SARS-CoV-2. Also, their positions on 3D structures are shown in Fig 3. Proposed epitope of B cell is conserved in all strains are shown here in the structural level of envelope protein of virus.

**Fig. 3.**
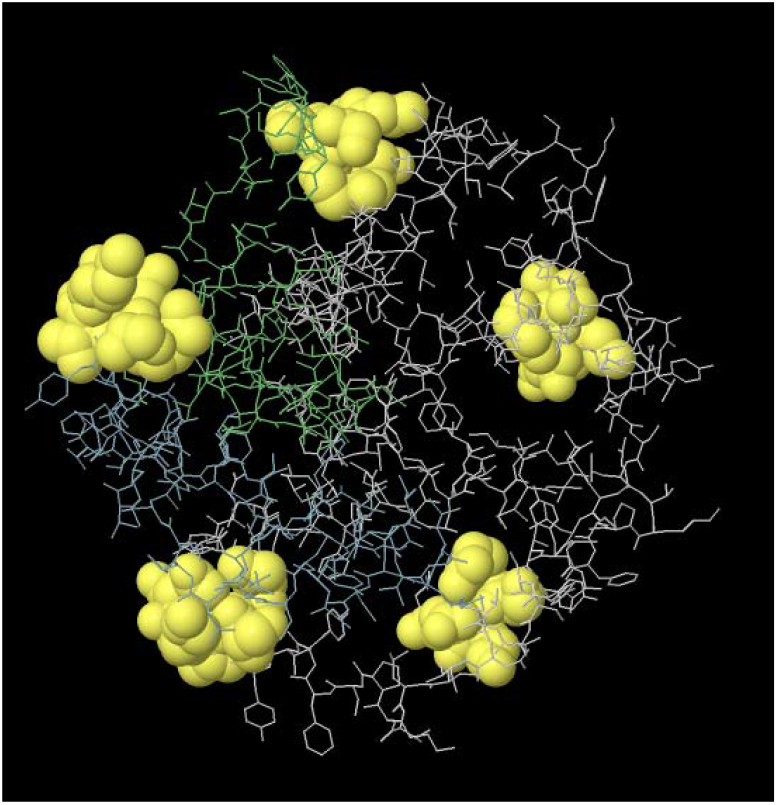
2D representation of conformational epitopes on five chains of E protein of SARS-CoV-2 showing the sequence composed of antigenic epitopes that will come in direct contact with immune receptor. The epitopes are represented by yellow surface, represent the surface Accessibility of SRVKNL, and the bulk of the E protein is represented in sticks.

**Table-1.** List of conformational B-cell epitopes of the E protein of SARS-CoV-2.

| No. | Residues | Number of residues | Score | Allergen | Antigenic |
| --- | --- | --- | --- | --- | --- |
| 1 | A:S60, A:R61, A:V62, A:K63, A:N64, A:L65 | 6 | 0.767 | No | Yes |
| 2 | A:E8, A:T9, A:G10, A:T11, A:L12, A:I13, A:V14, A:S16 | 8 | 0.739 | Yes | No |
| 3 | A:L51, A:V52, A:K53, A:S55, A:F56, A:Y59 | 6 | 0.658 | Yes | No |
| 4 | A:A32, A:I33, A:T35, A:A36, A:L37, A:R38, A:L39, A:C40, A:A41, A:Y42 | 10 | 0.61 | No | Yes |

Envelope protein was subjected to Bepipred linear epitope prediction, Emini surface accessibility, Karplus & Schulz Flexibility Prediction, Parker hydrophilicity and Chou and Fasman beta turn prediction methods in IEDB, that predict the probability of specific regions in the protein to bind to B cell receptor, being in the surface, being immunogenic, being in a hydrophilic region and being in a beta turn region, respectively, Fig.4.

**Fig. 4.**
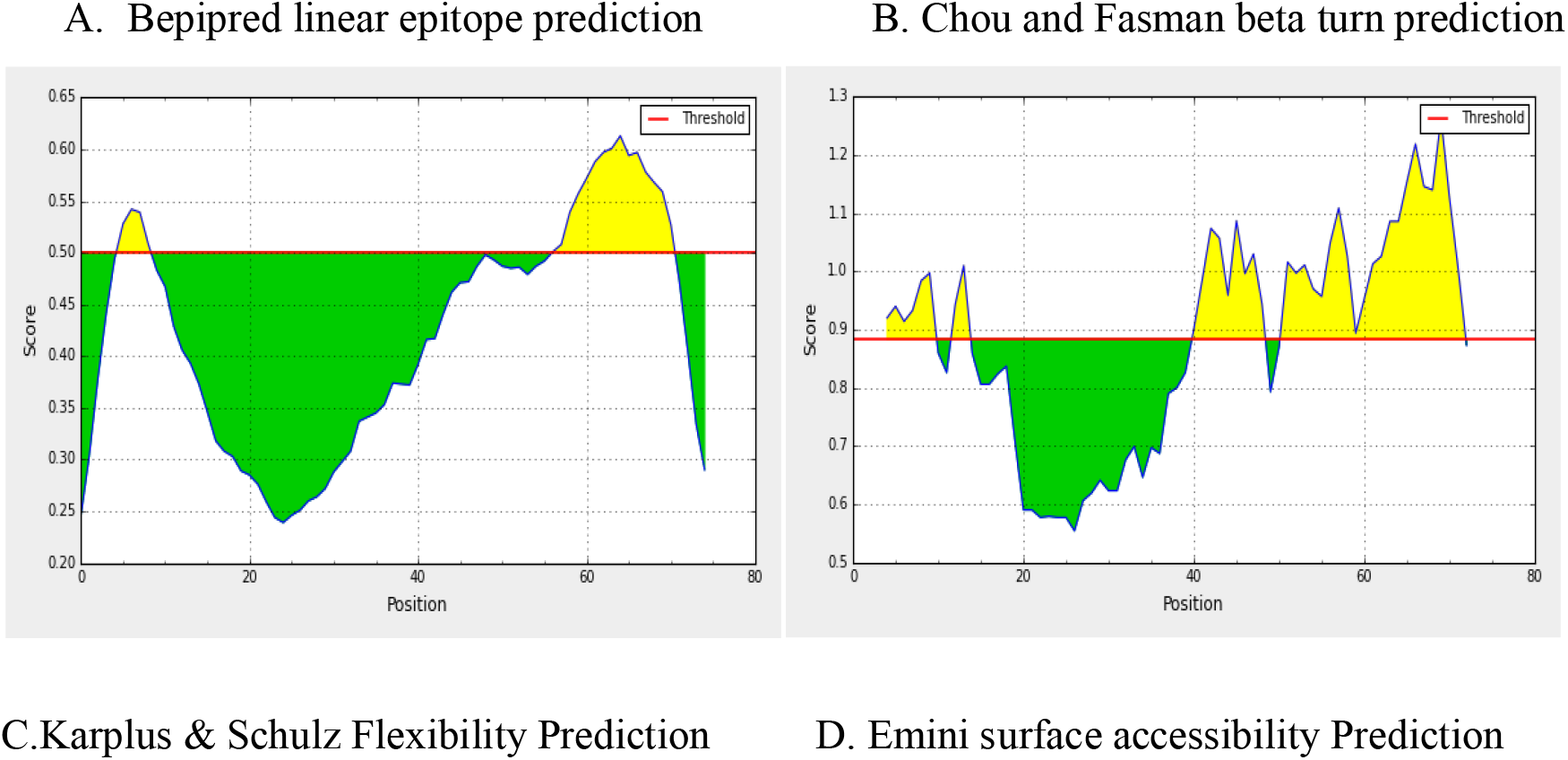
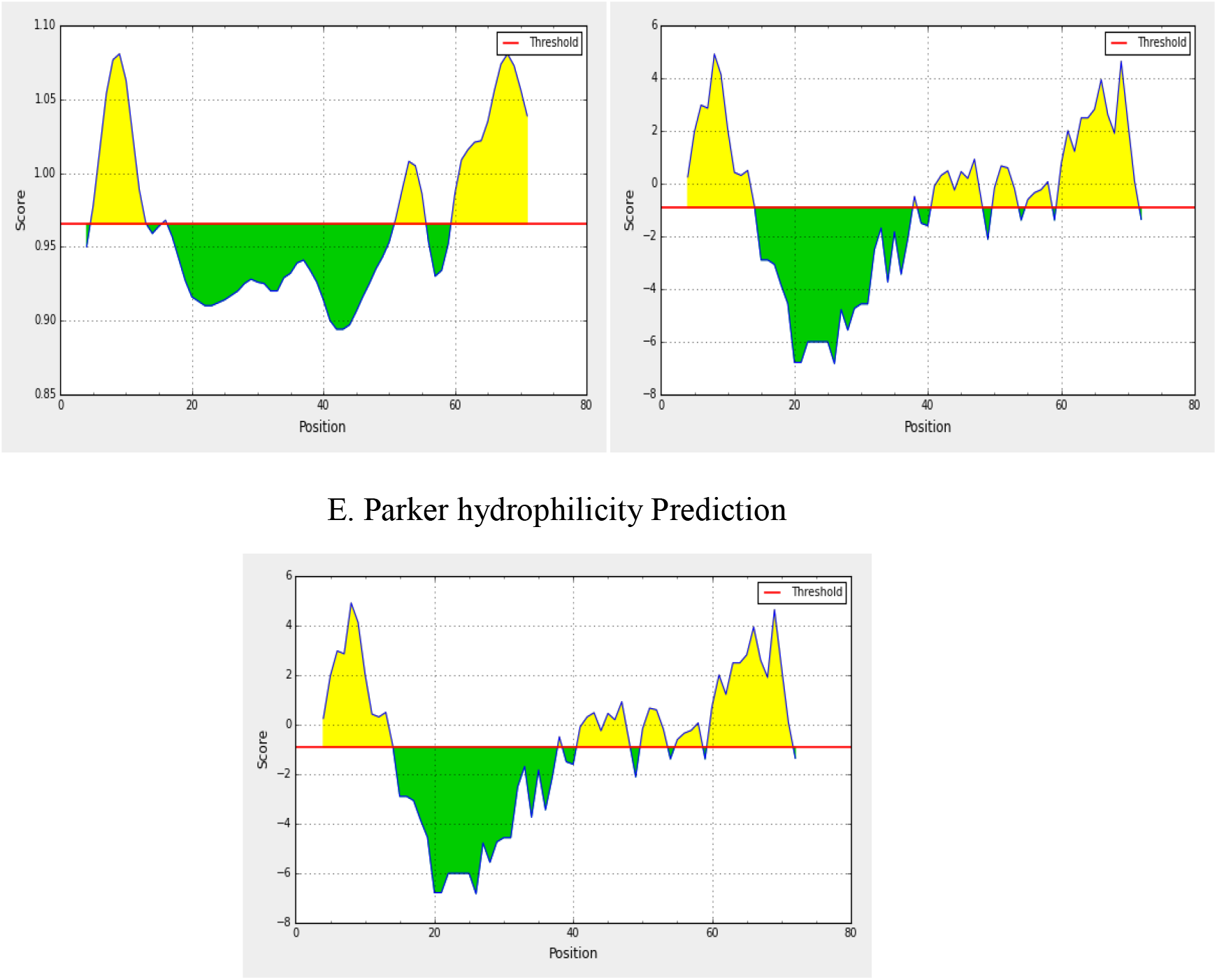
Prediction of B-cell epitopes by different scales/ parameters (A to E), yellow areas above threshold (red line) are proposed to be a part of B cell epitope. Epitope SRVKNL from 60-65 position satisfies threshold values of all the parameters.

In the Bepipred Linear Epitope Prediction method; the average binders score of envelope protein to B cell was 0.421, with a maximum of 0.613 and a minimum of - 0.239, all values equal or greater than the default threshold 0.023 were predicted to be a potential B cell binders. In Emini surface accessibility prediction; the average surface accessibility areas of the protein were scored as 1.000, with a maximum of 4.316 and a minimum of 0.088, all values equal or greater than the default threshold 1.0 were potentially in the surface. The default threshold of antigenicity of the protein was 1.119; with a maximum of 1.262 and a minimum of 0.947. In Parker’s hydrophilicity prediction; the average hydrophilicity score of the protein was 1.480, with a maximum of 4.929 and a minimum of −6.843, all values equal or greater than the default threshold −0.911 were potentially hydrophilic. The Chou and Fasman beta turn prediction method was used with the default threshold 0.883 with a maximum of 1.264 and a minimum of 0.883 for more confirmation for the prediction of the epitope to elicit B cell employed. The Karplus & Schulz Flexibility Prediction method was used with the default threshold 0.965 with a maximum of 1.081 and a minimum of 0.894 for more confirmation for the prediction of the epitope to elicit B cell employed. ^60^SRVKNL^65^ epitope was found to satisfy the threshold of all parameters/scales of B cell epitope prediction (Fig. 4).

### Prediction of Cytotoxic T-lymphocyte Epitopes and Interaction with MHC Class I

Envelope protein from the SARS-CoV-2 was analyzed using the IEDB MHC-1 binding prediction tool to predict the T cell epitope suggested interacting with different types of MHC Class I alleles. Based on NetCTL and SMM-based IEDB MHC-I binding prediction tools with higher affinity (IC50 less than 200) were predicted to interact with different MHC-1 alleles. The predicted total score of proteasome score, tap score, MHC score, processing score, and MHC-I binding are summarized as a total score in Table 2. These epitopes are antigenic and nonallergic. The peptide FLAFVVFLL from 20 to 29 (Fig. 5) had higher affinity to interact with 5 alleles (HLA-A*02:01, HLA-A*02:06, HLA-B*15:02, HLA-C*03:03, and HLA-A*68:02), followed by FLLVTLAIL from 26 to 34, and LLFLAFVVF from, 18 to 26 that had an affinity to interact with 5-6 alleles for each. The epitopes and their corresponding MHC-1 alleles are shown in Table 2. Among these 5 T-cell epitopes, 9-mer epitope, FLAFVVFLL was found to have the highest immunogenicity which was maximum than above said epitope and found to have more number of allelic interactions with good population coverage than other epitopes.

**Fig. 5.**
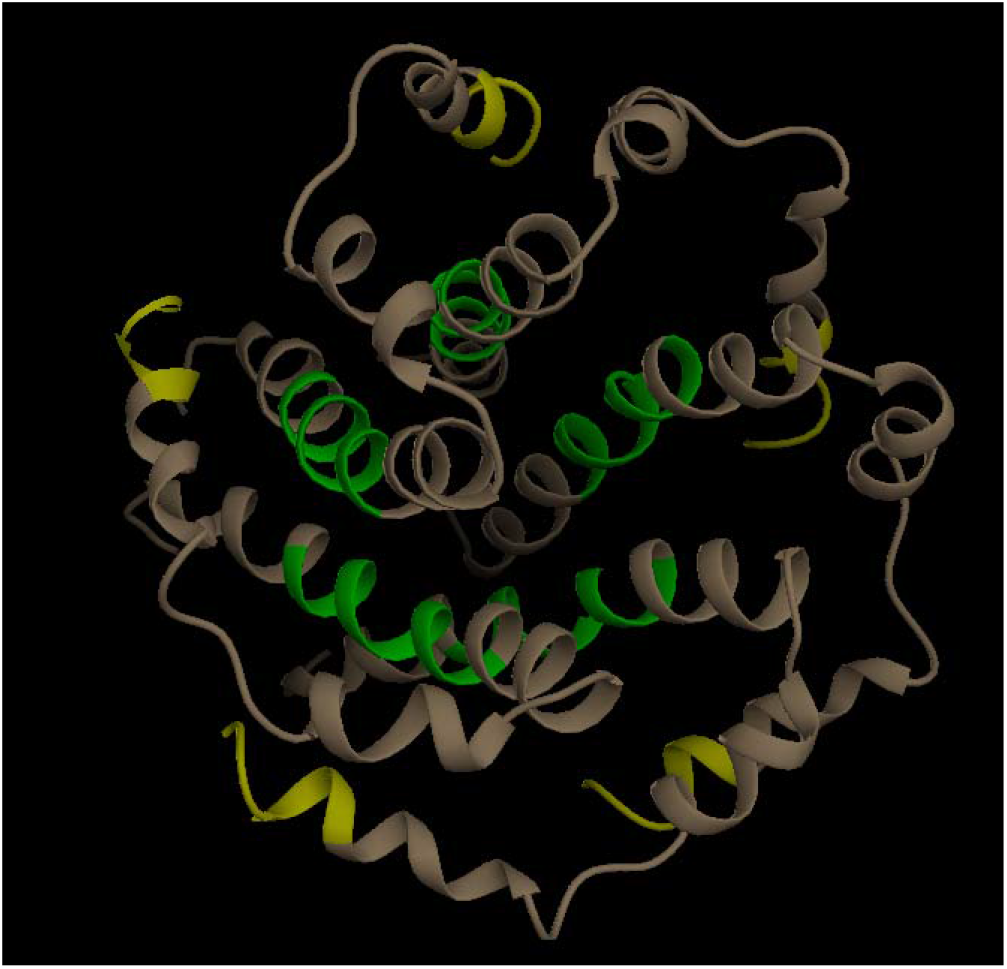
Proposed T-Cell epitopes, FLAFVVFL (green) and B cell epitopes, SRVKNL (yellow) in pantamer structure of E protein of SARS-CoV-2.

**Table-2.**
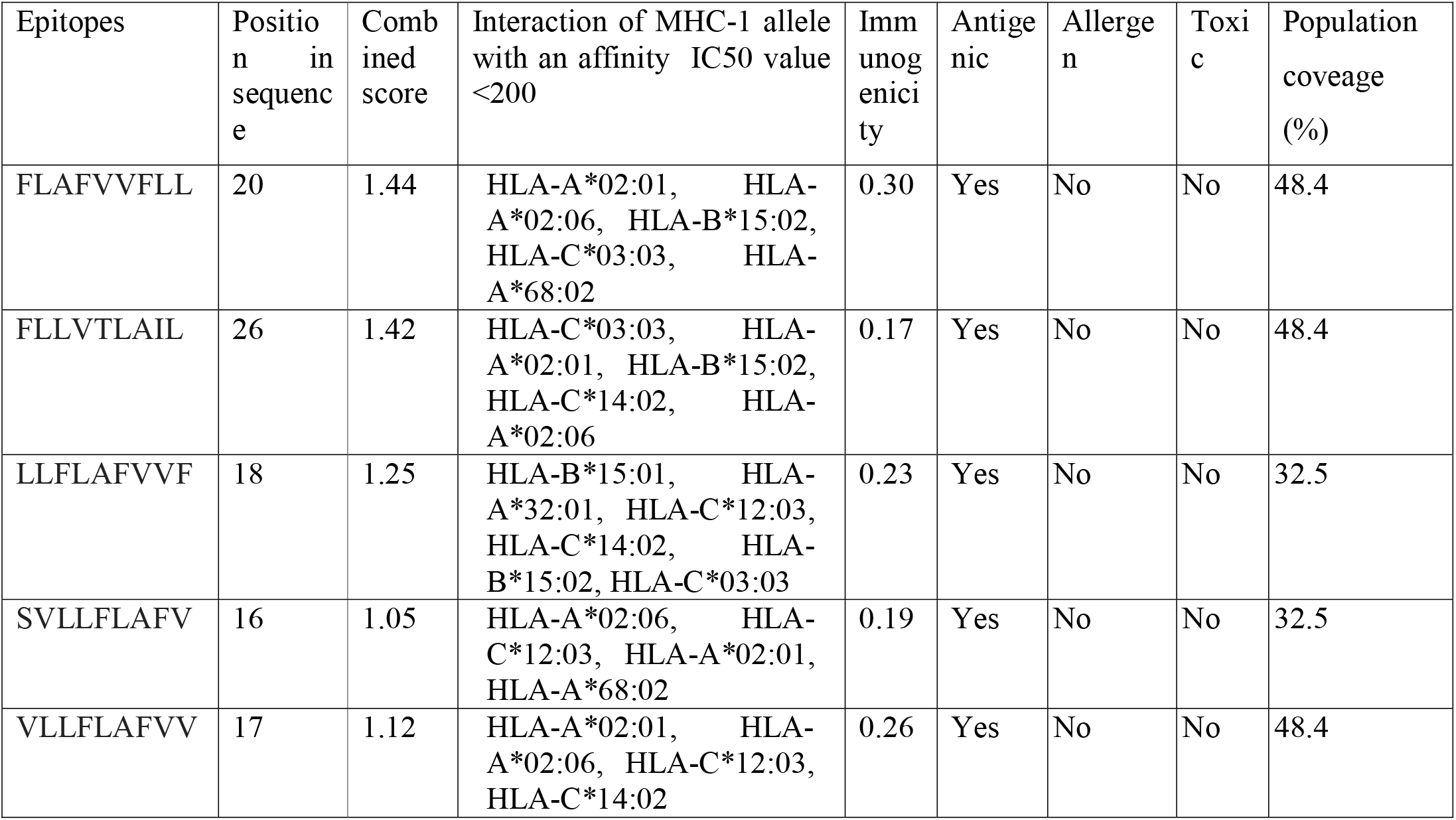
List of CTL epitopes that have good combined score, antigenicity, nonallergic, immunogenic, and binding with an affinity IC50 value less than 200 with the MHC I alleles.

### Prediction of Helper T-lymphocyte Epitopes and Interaction with MHC Class II

By the same way in IEDB MHC-1 binding prediction tool, T-cell epitopes from the SARS-CoV-2 were analyzed using the MHC-II binding prediction method; based on SMM based NetMHCIIpan with IC50 less than 200. There were top 5 predicted epitopes found to be nonallergic and antigenic interact with MHC-II alleles for which the peptide (core) FLAFVVFLL and LLVTLAILT had high affinity to interact with nine alleles (HLA-DRB1*04:05, HLA-DRB1*04:04, HLA-DRB1*15:01, HLA-DRB1*04:01, HLA-DRB1*01:01, HLA-DRB1*07:01, HLA-DRB1*04:04, HLA-DRB1*09:01, HLA-DRB5*01:01) and (HLA-DRB1*01:01, HLA-DRB1*15:01, HLA-DRB1*07:01, HLA-DRB1*04:04, HLA-DRB1*07:01, HLA-DRB1*04:05, HLA-DRB1*07:01, HLA-DRB1*15:01, HLA-DRB1*11:01) respectively. Moreover, the FLAFVVFLL epitope found to have maximum population coverage. The result is listed in Table 3. These epitopes conform that these epitopes have the capability to induce IFN-γ to show positive results.

**Table- 3.** List of HTL epitopes that are antigenic, nonallergic, and binding with an affinity IC50 value less than 200nm with MHC-II alleles.

| Epitope | Position in sequence | Interaction of MHC-II alleles having IC50 value < 200nm | Antigenic | Allergen | Toxicity | IFN-γ epitope prediction | Population coverage (%) |
| --- | --- | --- | --- | --- | --- | --- | --- |
| VLLFLAFVV | 16 | HLA-DRB1*15:01, HLA-DRB1*12:01, HLA-DRB1*04:05, HLA-DRB1*04:04, HLA-DRB1*01:01, HLA-DRB5*01:01, HLA-DRB1*07:01 | Yes | No | No | Positive | 53 |
| FLAFVVFLL | 20 | HLA-DRB1*04:05, HLA-DRB1*04:04, HLA-DRB1*15:01, HLA-DRB1*04:01, HLA-DRB1*01:01, HLA-DRB1*07:01, HLA-DRB1*04:04, HLA-DRB1*09:01, HLA-DRB5*01:01 | Yes | No | No | Positive | 62 |
| LLVTLAILT | 27 | HLA-DRB1*01:01, HLA-DRB1*15:01, HLA-DRB1*07:01, HLA-DRB1*04:04, HLA-DRB1*07:01, HLA-DRB1*04:05, HLA-DRB1*07:01, HLA-DRB1*15:01, HLA-DRB1*11:01 | Yes | No | No | Positive | 57 |
| LAILTALRL | 31 | HLA-DRB1*01:01, HLA-DRB1*15:01, HLA-DRB1*07:01, HLA-DRB1*04:04, HLA-DRB1*04:05, HLA-DRB1*11:01 | Yes | No | No | Positive | 57 |
| YVYSRVKNL | 57 | HLA-DRB1*04:01, HLA-DRB1*04:05, HLA-DRB1*07:01, HLA-DRB1*11:01, HLA-DRB5*01:01, HLA-DRB1*01:01 | Yes | No | No | Positive | 48 |

There were several overlapping between MHC class I epitopes and MHC class II epitopes. The overlapping epitopes are found from amino acid sequences 16 to 34 for MHC class I and II alleles, suggesting the possibility of antigen presentation to immune cells via both MHC class I and II pathways i.e. ^16^SVLLFLAFV^24^, ^26^FLLVTLAIL^34^, ^17^VLLFLAFVV^26^ and ^18^LLFLAFVVF^26^ (Table-2 and 3).

### Analysis of the Population Coverage

Epitopes that are suggested interacting with MHC-I and II alleles (especially high affinity binding epitopes and that can bind to a different set of alleles) were selected for population coverage analysis. The results of population coverage of all epitopes are listed in Table 2 and 3. FLAFVVFLL epitope that interacts with most frequent MHC class I and II alleles gave a high percentage against the whole world population by the IEDB population coverage tool. The maximum class I and II combined population coverage (84.88%) for this proposed epitope was found in North America (Table-4), while the higher population coverage in Europe (83.87%) and East Asia (81.61%) followed by South Asia (65.17%) and North Africa (65.31%) then Northeast Asia (64.8%) and Southeast Asia (61.1%). Table 4 represents the populations for which the MHC I and II Class Combined coverage of other areas.

**Table-4.**
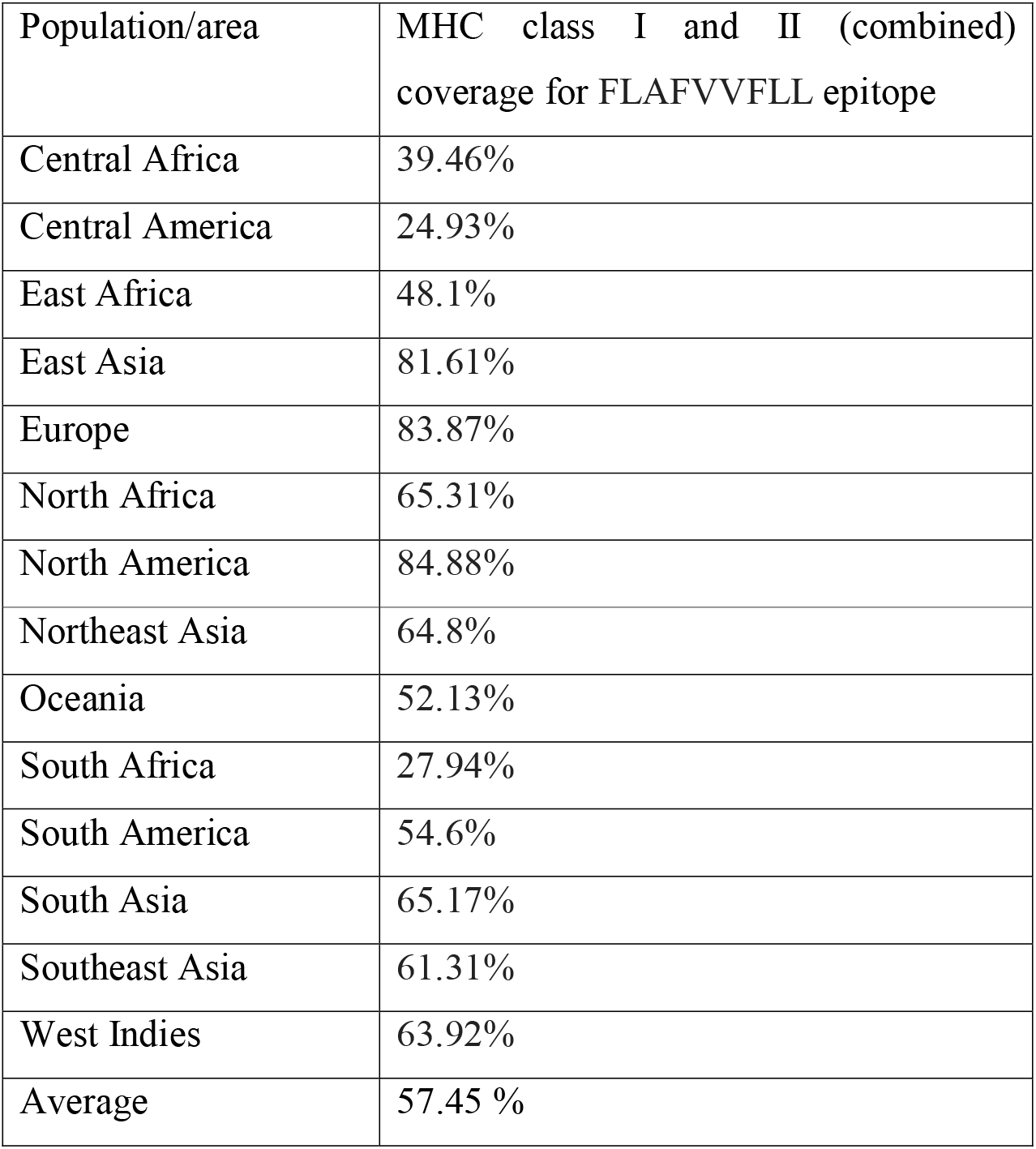
Population coverage of FLAFVVFLL epitope with MHC class I and II (combined) in the world.

### Molecular Docking of MHC I and II alleles with predicted T cell Epitope

The predicted T cell epitope FLAFVVFLL that interacted with selected human’s MHC-1and II alleles were used as ligands (Fig. 6) to detect their interaction with alleles /receptors, by docking techniques using on-line software Patchdock. After successful docking by PatchDock, the refinement and re-scoring of the docking results were carried out by the FireDock server. After refinement of the docking scores, the FireDock server generates global energies/ binding energies for the best solutions. Chimera was used to visualize the best results. The 3D structure of epitopes was predicted using PEP-FOLD and energy minimization was carried out by using Chimera. Based on the binding energy in kcal/mol unit, the lowest binding energy (kcal/mol) was selected to obtain a best binding (pose) and to predict real CTL and HTL epitope as possible.

**Fig. 6.**
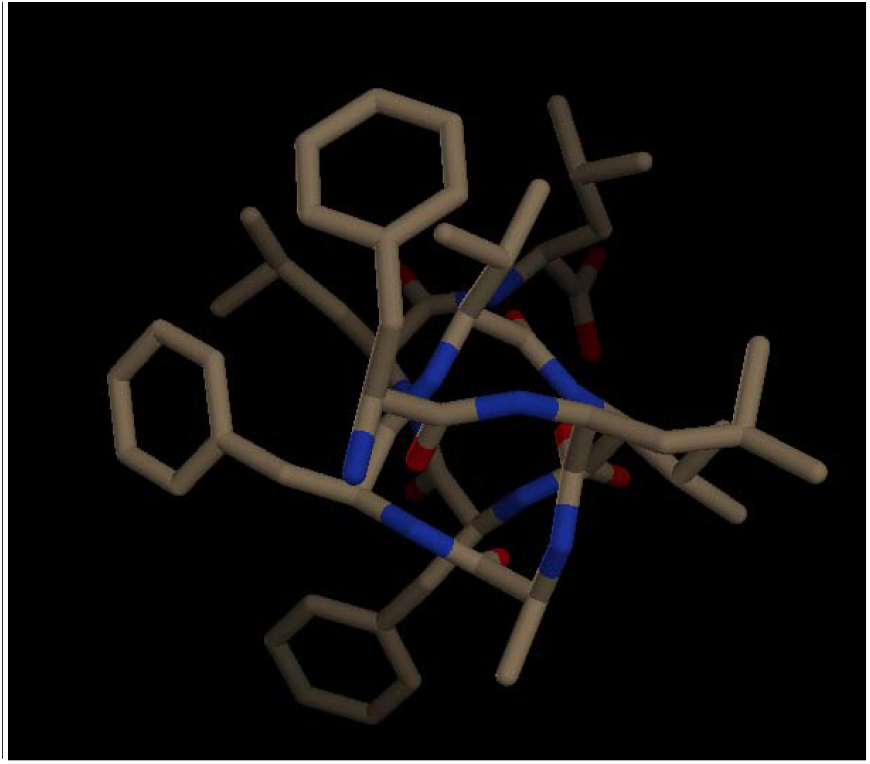
Peptide Structure of predicted epitope, FLAFVVFLL for MHC I and II prepared by PEP-FOLD.

The receptors used for docking studies included reported HLAs, HLA-C*03:03 (PDB ID: 1EFX) for class I and HLA-DRB1*01:01 (PDB ID: 1AQD) for class II. HLA-C*03:03 and HLA-DRB1*01:01 was observed to have the interaction with the FLAFVVFLL epitope with lower binding energy, −50.11 kCal/mol and −73.72 kCal/mol respectively (Fig. 7 and 8). The predicted peptide showed significant binding affinities with all HLAs. Also, the binding energy of the predicted epitopes were compared with the binding energy of the already experimentally verified peptides and found to be negative [16].

**Fig. 7.**
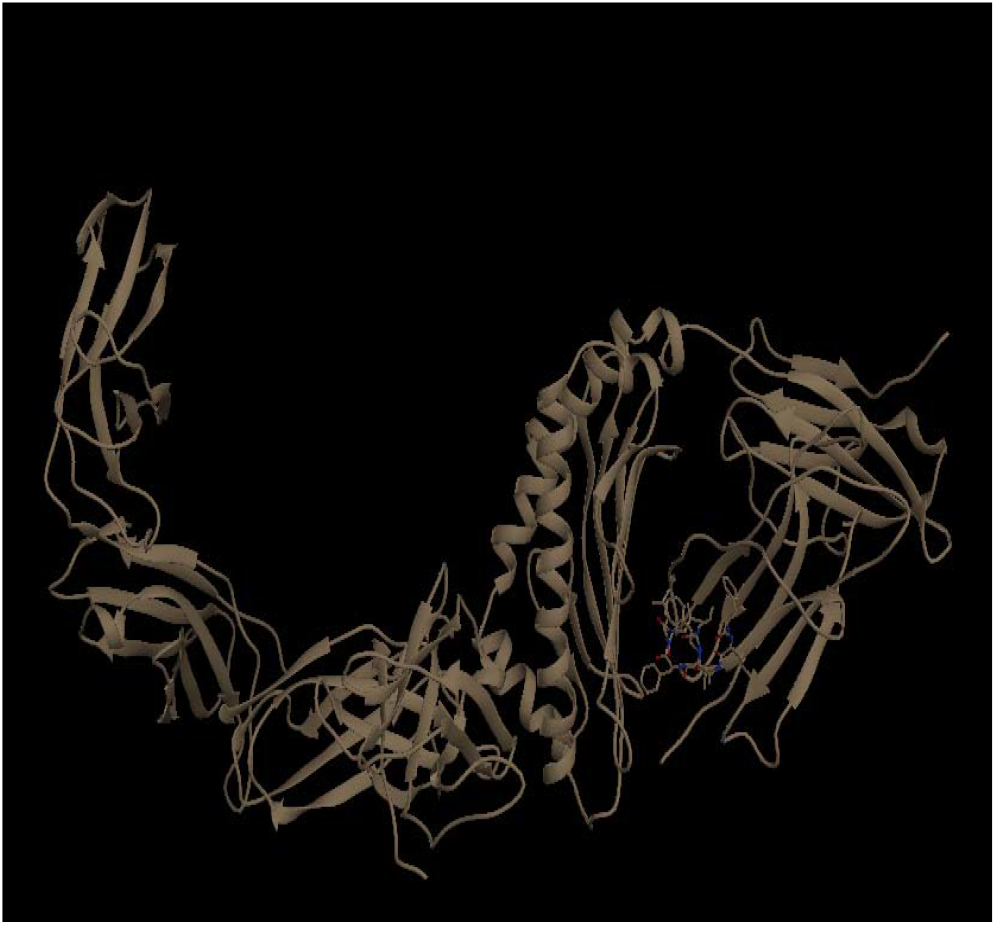
Docking sites of predicted peptide (FLAFVVFLL) against selected MHC I (1EFX) receptors.

**Fig. 8.**
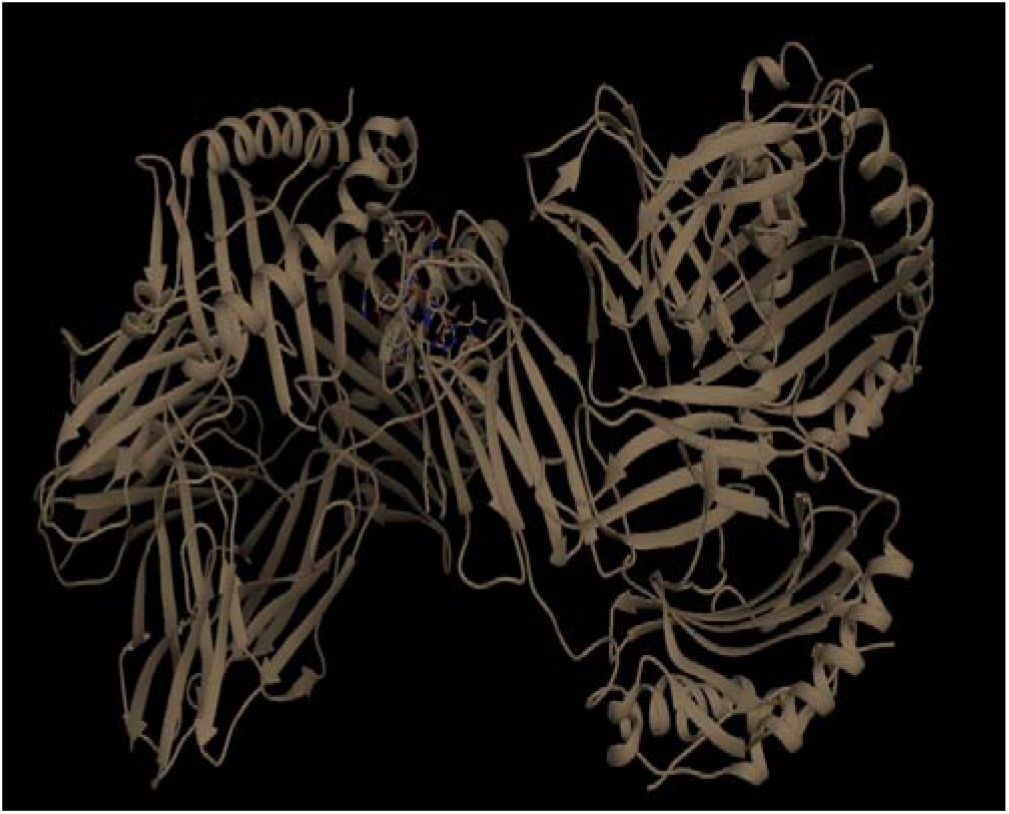
Docking sites of predicted peptide (FLAFVVFLL) against selected MHC II (1AQD) receptors.

## 4. Discussion

In this study, we aimed to determine the highly potential immunogenic epitopes for B and T cells, the prime molecules of humoral, and cell mediated immunity as peptide vaccine candidates for COVID-19 infection using the envelope protein as a target. Conservancy in E protein in the SARS-CoV-2 was found promising for peptide vaccine design. To determine potential and effective peptide antigen for B cell epitopes should get above threshold scores in Bepipred linear epitope prediction, Emini surface accessibility, Parker hydrophobicity, Karplus & Schulz Flexibility Prediction, and Chou and Fasman beta turn prediction methods in IEDB. SRVKNL epitope was found passing the thresholds of all prediction parameters in envelope protein. SRVKNL epitope was found to be antigenic, non-toxin, nonallergic and conserved in all sequences of SARS-CoV-2 consider in this study (supplementary file). The conformational epitopes were predicted by using a 3D structure of the E protein. The B-cell epitopes residues, SRVKNL located on the surface of the E protein had good Protrusion Index (PI) score (0.76) were indicative of high accessibility. Ellipsoid value of PI 0.76 indicates that 76% of protein residues lie within ellipsoid and the remaining 24% residues lie outside. PI score and solvent accessibility are directly proportional to each other if the PI score is higher; maximum is the solvent accessibility of the residues [50]. Thus, this epitope enables direct interactions with an immune receptor, which could be the putative vaccine candidates.

Since the immune response of T cell is long lasting response comparing with B cell, where the antigen can easily escape the antibody memory response [18] additionally, CD8+ T and CD4+ T cell responses play a major role in antiviral immunity [16], designing of a vaccine against T cell epitope is much more promising. FLAFVVFLL epitope could be used as a potential candidate because it had a maximum combined score and immunogenic score. Moreover, it possessed the maximum number of HLA binding alleles amongst other CTL and HTL. This epitope was found to be antigenic, non-toxin and nonallergic. An ideal epitope should be highly conserved. The conservancy analysis of this epitopes indicated that this epitope was found to have been conserved in all sequences of the SARS-CoV-2 consider in this study. This consistency of immunological features of epitopes indicates that these parameters fulfill all criteria of further screening. Further, docking study was performed with FLAFVVFLL epitope to check interaction with MHC class I and II alleles. The binding affinity of the epitope, FNLTLLEPV with MHC I and II alleles shows a very high binding affinity with negative binding energy. This epitope alone showed 57% coverage in the world, a maximum population coverage found in North America (85 %) and in East Asia (82%).

Also, the 9-mer CTL epitopes (^16^SVLLFLAFV^24^, ^26^FLLVTLAIL^34^, ^17^VLLFLAFVV^26^ and ^18^LLFLAFVVF^26^) from 16 to 34 amino acid sequence of E protein were able to interact with MHC class I alleles and could become universal peptide based vaccine against COVID-19. We found these CTL epitopes to be HTL epitopes. The overlapping between MHC Class I and II T cell epitopes suggested the possibility of antigen presentation to immune cells via both MHC class I and II pathways especially the overlapping sequences.

To conclude, by using E protein one epitope, SRVKNL was proposed for an international therapeutic peptide vaccine for B cell. Regarding T cell, the FLAFVVFLL epitope was highly recommended as a therapeutic peptide vaccine to interact with both MHC class I and II. We recommend *in vitro* and *in vivo* validation for the efficacy and efficiency of these predicted candidate epitopes as a vaccine as well as to be used as a diagnostic screening test.

## Financial support & sponsorship

None

## Conflict of interest

None

## Author contribution

Renu Jakhar conducted the study, performed *in silico* analysis and wrote the manuscript. S.K. Gakhar plans the study and revises the manuscript.

## Notes

### Competing Interest Statement

The authors have declared no competing interest.

## References

1. Lu H, Stratton CW, Tang YW. Outbreak of Pneumonia of Unknown Etiology in Wuhan China: the Mystery and the Miracle. Journal of Medical Virology 2020.

2. Wu F, Zhao S, Yu B, Chen YM, Wang W, Song ZG et al. A new coronavirus associated with human respiratory disease in China. Nature 2020; 3:1–5.

3. Benvenuto D, Giovannetti M, Ciccozzi A, Spoto S, Angeletti S, Ciccozzi M. The 2019-new coronavirus epidemic: evidence for virus evolution. Journal of Medical Virology 2020.

4. Chen Y, Liu Q, Guo D. Emerging coronaviruses: genome structure, replication, and pathogenesis. Journal of medical virology 2020.

5. Chan JF, Yuan S, Kok KH, To KK, Chu H, Yang J et al. A familial cluster of pneumonia associated with the 2019 novel coronavirus indicating person-to-person transmission: a study of a family cluster. The Lancet 2020; 15;395(10223):514–23.

6. Corman VM, Landt O, Kaiser M, Molenkamp R, Meijer A, Chu DK et al. Detection of 2019 novel coronavirus (2019-nCoV) by real-time RT-PCR. Eurosurveillance 2020; 25.

7. Ji W, Wang W, Zhao X, Zai J, Li X. Cross-species transmission of the newly identified coronavirus 2019-nCoV. Journal of Medical Virology 2020; 92(4):433–40.

8. Zhang N, Wang L, Deng X, Liang R, Su M, He C et al. Recent advances in the detection of respiratory virus infection in humans. Journal of Medical Virology 2020.

9. Hui DS, Azhar E, Madani TA, Ntoumi F, Kock R, Dar O et al. The continuing 2019-nCoV epidemic threat of novel coronaviruses to global health—The latest 2019 novel coronavirus outbreak in Wuhan, China. International Journal of Infectious Diseases 2020; 91:264–6.

10. Shi J, Zhang J, Li S, Sun J, Teng Y, Wu M. Epitope-based vaccine target screening against highly pathogenic MERS-CoV: an in silico approach applied to emerging infectious diseases. PloS one 2015; 10:e0144475.

11. Srivastava S, Kamthania M, Singh S, Saxena AK, Sharma N. Structural basis of development of multi-epitope vaccine against middle east respiratory syndrome using in silico approach. Infection and drug resistance 2018;11:2377

12. Qamar MT, Saleem S, Ashfaq UA, Bari A, Anwar F, Alqahtani S. Epitope-based peptide vaccine design and target site depiction against Middle East Respiratory Syndrome Coronavirus: an immune-informatics study. Journal of translational medicine 2019; 17:362.

13. Yong CY, Ong HK, Yeap SK, Ho KL, Tan WS. Recent Advances in the Vaccine Development Against Middle East Respiratory Syndrome-Coronavirus. Frontiers in microbiology 2019; 10:1781.

14. Ahmed SF, Quadeer AA, McKay MR. Preliminary identification of potential vaccine targets for the COVID-19 coronavirus (SARS-CoV-2) based on SARS-CoV immunological studies. Viruses 2020; 12:254.

15. Schoeman D, Fielding BC. Coronavirus envelope protein: current knowledge. Virology journal 2019; 16:69.

16. Liu J, Sun Y, Qi J, Chu F, Wu H, Gao F et al. The membrane protein of severe acute respiratory syndrome coronavirus acts as a dominant immunogen revealed by a clustering region of novel functionally and structurally defined cytotoxic T-lymphocyte epitopes. The Journal of infectious diseases 2010; 202:1171–80.

17. Grifoni A, Sidney J, Zhang Y, Scheuermann RH, Peters B, Sette A. A Sequence Homology and Bioinformatic Approach Can Predict Candidate Targets for Immune Responses to SARS-CoV-2. Cell host & microbe 2020.

18. Xie Q, He X, Yang F, Liu X, Li Y, Liu Y et al. Analysis of the genome sequence and prediction of B-cell epitopes of the envelope protein of Middle East respiratory syndrome-coronavirus. IEEE/ACM transactions on computational biology and bioinformatics 2017; 15:1344–50.

19. Wang Y, Liu L. The membrane protein of severe acute respiratory syndrome coronavirus functions as a novel cytosolic pathogen-associated molecular pattern to promote beta interferon induction via a Toll-like-receptor-related TRAF3-independent mechanism. MBio 2016; 7:e01872–15.

20. Ying T, Du L, Ju TW, Prabakaran P, Lau CC, Lu L, Liu et al. Exceptionally potent neutralization of Middle East respiratory syndrome coronavirus by human monoclonal antibodies. Journal of virology 2014; 88:7796–805.

21. Wang L, Shi W, Joyce MG, Modjarrad K, Zhang Y, Leung K et al. Evaluation of candidate vaccine approaches for MERS-CoV. Nature communications 2015; 6:1–1.

22. Rappuoli R. Reverse vaccinology. Curr Opin Microbiol 2000; 3:445–50.

23. Graham RL, Donaldson EF, Baric RS. A decade after SARS: strategies for controlling emerging coronaviruses. Nat Rev Microbiol 2013 D; 11:836–48.

24. Purcell AW, McCluskey J, Rossjohn J. More than one reason to rethink the use of peptides in vaccine design. Nat Rev Drug Discov 2007; 6:404–14.

25. Doytchinova IA, Flower DR. VaxiJen: a server for prediction of protective antigens, tumour antigens and subunit vaccines. BMC Bioinformatics 2007; 8.

26. Waterhouse A, Bertoni M, Bienert S, Studer G, Tauriello G, Gumienny R et al. SWISS-MODEL: homology modelling of protein structures and complexes. Nucleic acids research 2018; 46: W1: W296–W303.

27. Pettersen EF, Goddard TD, Huang CC, Couch GS, Greenblatt DM, Meng EC et al. UCSF Chimera, a visualization system for exploratory research and analysis. Journal of computational chemistry 2004; 25:1605–1612.

28. Ramachandran GN, Ramakrishnan C, Sasisekharan V. Stereochemistry of polypeptide chain configurations. J Mol Biol 1963; 7:95–9.

29. Andersen PH, Nielsen M and Lund O. Prediction of residues in discontinuous B-cell epitopes using protein 3D structures. Protein Sci 2006; 15:2558–67.

30. Emini EA, Hughes JV, Perlow DS, Boger J. Induction of hepatitis A virus-neutralizing antibody by a virus-specific synthetic peptide. J Virol 1985; 55:836–9.

31. Kolaskar AS, Tongaonkar PC. A semi-empirical method for prediction of antigenic determinants on protein antigens. FEBS Lett 1990; 276:172–4.

32. Parker JM, Guo D, Hodges RS. New hydrophilicity scale derived from high-performance liquid chromatography peptide retention data: correlation of predicted surface residues with antigenicity and X ray-derived accessible sites. Biochemistry 1986; 25:5425–32.

33. Chou PY, Fasman GD. Prediction of the secondary structure of proteins from their amino acid sequence. Adv Enzymol Relat Areas Mol Biol 1978; 47:45–148

34. Karplus PA, Schulz GE. Prediction of chain flexibility in proteins. Naturwissenschaften 1985; 72:212–3.

35. Wilson JA, Hart MK. Protection from Ebola virus mediated by cytotoxic T lymphocytes specific for the viral nucleoprotein. J Virol 2001; 75:2660–4.

36. Dimitrov I, Flower DR, Doytchinova I. AllerTOP - a server for in silico prediction of allergens. BMC Bioinformatic 2013; 14: Suppl 6:S4.

37. Gupta S, Kapoor P, Chaudhary K, Gautam A, Kumar R, Raghava GP. Open Source Drug Discovery Consortium. In silico approach for predicting toxicity of peptides and proteins. PloS one 2013; 8(9).

38. Larsen MV, Lundegaard C, Lamberth K, Buus S, Lund O, Nielsen M. Large-scale validation of methods for cytotoxic T-lymphocyte epitope prediction. BMC bioinformatics 2007; 8:424.

39. Buus S, Lauemoller SL, Worning P, Kesmir C, Frimurer T, Corbet S et al. Sensitive quantitative predictions of peptide-MHC binding by a ‘Query by Committee’artificial neural network approach. Tissue antigens 2003; 62:378–84.

40. Vita R, Overton JA, Greenbaum JA, Ponomarenko J, Clark JD, Cantrell JR et al. The immune epitope database (IEDB) 3.0. Nucleic acids research 2015; 43:D405–12.

41. Calis JJ, Maybeno M, Greenbaum JA, Weiskopf D, De Silva AD, Sette A et al. Properties of MHC class I presented peptides that enhance immunogenicity. PLoS computational biology 2013; 10.

42. Karosiene E, Rasmussen M, Blicher T, Lund O, Buus S, Nielsen M. NetMHCIIpan-3.0, a common panspecific MHC class II prediction method including all three human MHC class II isotypes, HLA-DR,HLA-DP and HLA-DQ. Immunogenetics 2013; 65:711–24.

43. Wilson JA, Hart MK. Protection from Ebola virus mediated by cytotoxic T lymphocytes specific for the viral nucleoprotein. J Virol 2001; 75:2660–4.

44. Zhang L, Udaka K, Mamitsuka H, Zhu S. Toward more accurate pan-specific MHC-peptide binding prediction: a review of current methods and tools. Brief Bioinform 2012; 13:350–64.

45. Pandey R K, Bhatt T K, Prajapati V K. Novel immunoinformatics approaches to design multi-epitope subunit vaccine for malaria by investigating anopheles salivary protein. Scientific reports 2017; 8:1125.

46. Bui HH, Sidney J, Dinh K, Southwood S, Newman MJ, Sette A. Predicting population coverage of T-cell epitope-based diagnostics and vaccines. BMC Bioinformatics 2006; 7:153.

47. Thevenet P, Shen Y, Maupetit J, Guyon F, Derreumaux P, Tufféry P. PEP-FOLD: an updated de novo structure prediction server for both linear and disulfide bonded cyclic peptides. Nucleic Acids Res 2012; 40: W288–W293.

48. Deshpande N, Addess KJ, Bluhm WF, Merino-Ott JC, Townsend-Merino W, Zhang Q et al. The RCSB Protein Data Bank: a redesigned query system and relational database based on the mmCIF schema. Nucleic acids research 2005; 33:D233–7.

49. Schneidman-Duhovny D, Inbar Y, Nussinov R, Wolfson HJ. PatchDock and SymmDock: servers for rigid and symmetric docking. Nucleic Acids Res 2005; 33:W363–W367.

50. Jakhar R, Kumar P, Sehrawat N, Gakhar SK. A comprehensive analysis of amino-peptidase N1 protein (APN) from Anopheles culicifacies for epitope design using Immuno-informatics models. Bioinformation 2019; 15:600.

